# High nitrogen deposition is associated with phosphorus-efficient ectomycorrhizas in Europe’s Scots pine forests

**DOI:** 10.64898/2026.06.04.730184

**Authors:** Pierre Herinckx, Guillaume Delhaye, Martin I. Bidartondo, Roberta Gargiulo, Emil Ghaffar, Celestine Ruhmann, Max Ticehurst, Christopher Andrews, Vladislav Apuhtin, Caitlin Lewis, Päivi Merilä, Elena Vanguelova, Arne Verstraeten, Janna Wambsganss, Thomas Drouet, Laura M. Suz

## Abstract

Atmospheric inorganic nitrogen (N) deposition has been linked to increased tree phosphorus (P) deficiency and shifts in ectomycorrhizal (ECM) fungal community composition across Europe, but the underlying mechanisms remain poorly understood due to the scarcity of species-level studies of fungal physiology at large spatial scales. Here, we characterized ECM communities in nine Scots pine (*Pinus sylvestris* L.) stands across Europe’s largest N deposition gradient to gain mechanistic insight into N-driven ECM community shifts, by assessing morpho-physiological traits (i.e. soil exploration types and ECM root-tip level exoenzyme activities involved in organic N and P acquisition) on individual ectomycorrhizas. Our data revealed high functional variation in foraging strategies across species and sites, including within dominant ECM genera (*Cortinarius, Elaphomyces, Lactarius, Piloderma, Russula*). Shifts in community-level exoenzyme activities along the N deposition gradient were consistent with increasing P limitation, with a buffering effect of phosphomonoesterase activity on host nutritional status (i.e. reduced foliar N:P). These trends were mainly driven by interspecific differences in enzymatic profiles and species turnover along the gradient, rather than intraspecific variation within widespread species. Dominant low-biomass species in high N sites (e.g. *E. citrinopapillatus, L. subdulcis, R. ochroleuca*) were efficient P-foragers, with some displaying high oxidative activity, potentially hampering soil carbon storage under elevated N loads. These findings highlight the role of ECM species-specific traits in mediating ecosystem processes and can help understand the future of pine forests under chronic N pollution, with potential implications for applied forestry.

## Introduction

Tree nutritional status is degrading in Europe [1,2], which could have cascading effects on ecosystem services such as carbon (C) storage, nutrient cycling and tree health [3,4]. Atmospheric inorganic nitrogen (N) deposition, primarily as ammonium (NH_4_^+^) and nitrate (NO_3_^−^) derived from anthropogenic activities [5], is linked to imbalanced tree foliar N:P, reflecting an emerging N-induced phosphorus (P) limitation for forest productivity in Europe [1,6,7].

Ectomycorrhizal (ECM) fungi carry out tree nutrient uptake in most boreo-temperate forests, where they regulate major processes such as nutrient cycling and soil C storage [8–10]. Based on ECM community changes along environmental gradients in European forests, it has been proposed that increasing atmospheric inorganic N deposition causes steep changes in ECM diversity and community composition, notably towards a dominance of competitive nitrophilic taxa [11–13], which could impact tree nutrition and growth [14], and lead to a tipping point in forest dynamics [15]. Consequently, it was suggested that European critical loads for atmospheric N deposition should be revised downward from 10-20 kg N ha⁻¹ yr⁻¹ [16] to 5.8 kg N ha⁻¹ yr⁻¹ [13] in coniferous forests where many specialist ECM fungi are associated with a stronger decline in response to increased N availability [13,17]. In particular, nitrophilic fungi (e.g. *Russula ochroleuca, Lactarius subdulcis*) may outcompete C-costly nitrophobic ones (e.g. *Piloderma olivaceum, Cortinarius* spp.) under N-induced reduced host C allocation to root symbionts [18], resulting from lower investment in both extraradical mycelium production and specialization in organic N uptake [12,13,19].

However, despite a well-established picture of how ECM communities change in response to N deposition, the underlying mechanisms and their consequences on host tree and forest nutrient cycles remain poorly understood [20,21]. This is notably due to the scarcity of species-level studies of ECM fungal physiology at large spatial scales. Recent genomic insights show that ECM lineages retain degradative capacities inherited from their saprotrophic ancestors [22–24], potentially providing host trees with unavailable organic forms of N and P through the production of cell wall-bound exoenzymes, which substantially contribute to soil organic matter (SOM) transformation and cycling [25,26]. Along with nutrient-acquisition enzymes that directly hydrolyse organic sources of N (e.g. amino acids and peptides) and P (e.g. phospholipids and nucleic acids), some ECM fungi possess elevated lignocellulolytic capacities that enhance nutrient availability through the degradation of plant and fungal necromass [27]. These include oxidases, such as laccases and lignin peroxidases, targeting complex polyphenolic compounds [28,29], placing ECM physiological variation at the centre of soil C dynamics [30,31].

Despite recent findings linking gene copy numbers in ECM fungi with environmental variables and ecosystem processes [14], studies reveal that gene number is not a sufficient proxy for inferring microbial [32] and fungal [33,34] degradative activity. Hence, *in situ* measurement of physiological traits provides a direct measure of the interplay between genetic background and environment, and therefore allows a clearer understanding of the functional mechanisms leading to observed communities [35,36]. While previous studies have shown marked species-level differences in enzyme production among ECM fungi in relation to environmental factors such as host identity [37], soil chemistry [38], drought [39] or disturbance [40], large-scale studies using root-tip-level enzymatic traits along continuous gradients of atmospheric N deposition are critically needed to link ECM community composition to forest nutrient status. A key unresolved question is whether the increased abundance of nitrophilic species associated with chronic N deposition reflects enhanced P-acquisition capacities buffering the rise in tree foliar N:P [41]. Experimental results indicate that ECM hosts can regulate root colonization to favour better N-provisioning fungal species [42,43], but comparable evidence for host selection based specifically on P is still lacking [41]. The increasingly recognized role of ECM community composition and species-specific effects on ecosystem-level processes, such as forest growth rate and productivity [14], suggests that nitrophilic ECM fungi might be selected if they alleviate N-induced P limitation. In addition, the lack of ECM physiological data collected along continuous broad-scale gradients has long limited understanding of the contribution of intraspecific variation in shaping the distribution of widespread taxa, and its relative ecosystem-level implications, compared to well-documented elevated ECM species turnover in response to environmental changes [44–46].

Here, we present a rare investigation of the mechanisms driving changes in ECM community composition in response to N pollution across Europe’s Scots pine (*Pinus sylvestris* L.) forests, a common and widespread coniferous ECM tree of high economic value. We characterized the enzymatic profiles of nine ECM communities by measuring five root-associated exoenzyme activities involved in N and P acquisition from SOM. We asked: 1) How does the dominant nutrient acquisition strategy change along an atmospheric N deposition gradient? 2) Is there any intraspecific variation in enzymatic production and does it follow similar trends to the community? 3) Is there an effect of the dominant nutrient acquisition strategy on tree nutrient ratios (foliar N:P)? We hypothesized that:

i) Variation in community-level enzymatic profiles along the gradient mirrors the shift in nutrient limitation from N to P, consistent with optimal allocation theory [47] (H1).
ii) Community-level changes will be primarily driven by interspecific differences in trait values and species turnover along the gradient, with only minor intraspecific physiological variation in widespread fungi (H2). We expect elevated species replacement along the gradient to reflect an increase in efficient P-foraging ECM fungi, including through enhanced lignocellulolytic activity, at the expense of N-foragers at the high N end of the gradient.
iii) The ECM community functional response will have a positive buffering effect on tree nutritional balance (i.e. reduced foliar N:P), including through enhanced SOM oxidation (H3).

## Materials and Methods

### Selection of study plots and environmental variables

Nine monodominant Scots pine (*Pinus sylvestris* L.) stands (0.25 ha) were selected across six countries to encompass the largest possible gradient of total atmospheric inorganic N (sum of NH_4_^+^- and NO_3_^−^-N) deposition in Europe, based on previous data [13] (**Fig. 1**; **Table 1**). Eight plots are intensively monitored Level II sites of the International Co-operative Programme on Assessment and Monitoring of Air Pollution Effects on Forests (ICP Forests) [48], and one site is part of Scotland’s Sites of Special Scientific Interest (SSSIs). Both monitoring networks provided the *in situ* environmental covariates used in this study. The inorganic N deposition data are averaged site-specific measurements over six to eight years, and correspond to wet throughfall deposition, except for the Scottish site where only bulk wet deposition was available. Tree foliar N:P ratios were measured and calculated for three sites lacking these measurements (**Table 1**). Foliar N was quantified by combustion using the Dumas method and foliar P was extracted by acid digestion and quantified by Inductively Coupled Plasma Optical Emission Spectroscopy at NRM (Bracknell, UK). The four most N-polluted sites had foliar N:P ratios beyond the healthy range of 7.4 - 14.1 for *P. sylvestris* in Europe [49].

**Fig. 1.**
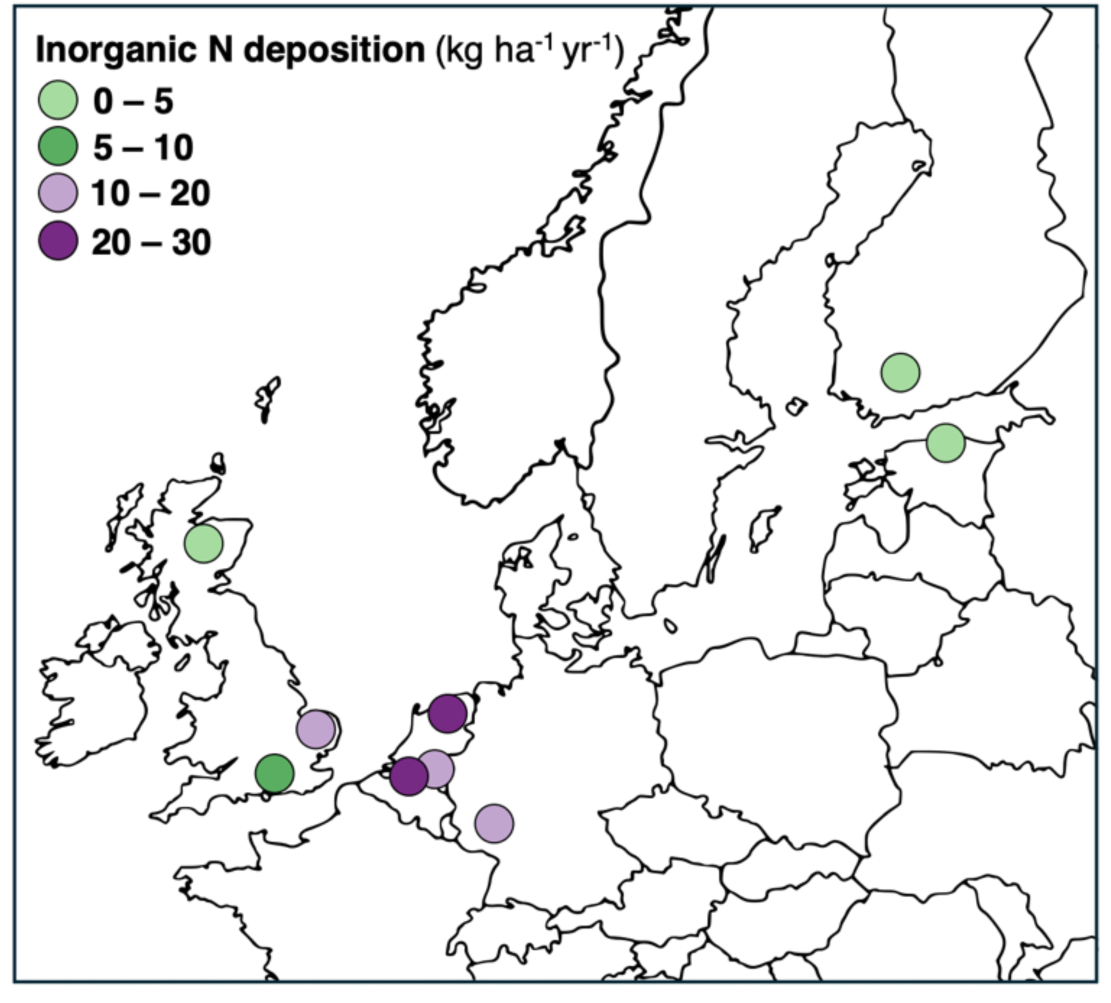
Location of the pine forest sampling plots in Northern Europe, coloured by level of atmospheric inorganic N deposition.

**Table 1.**
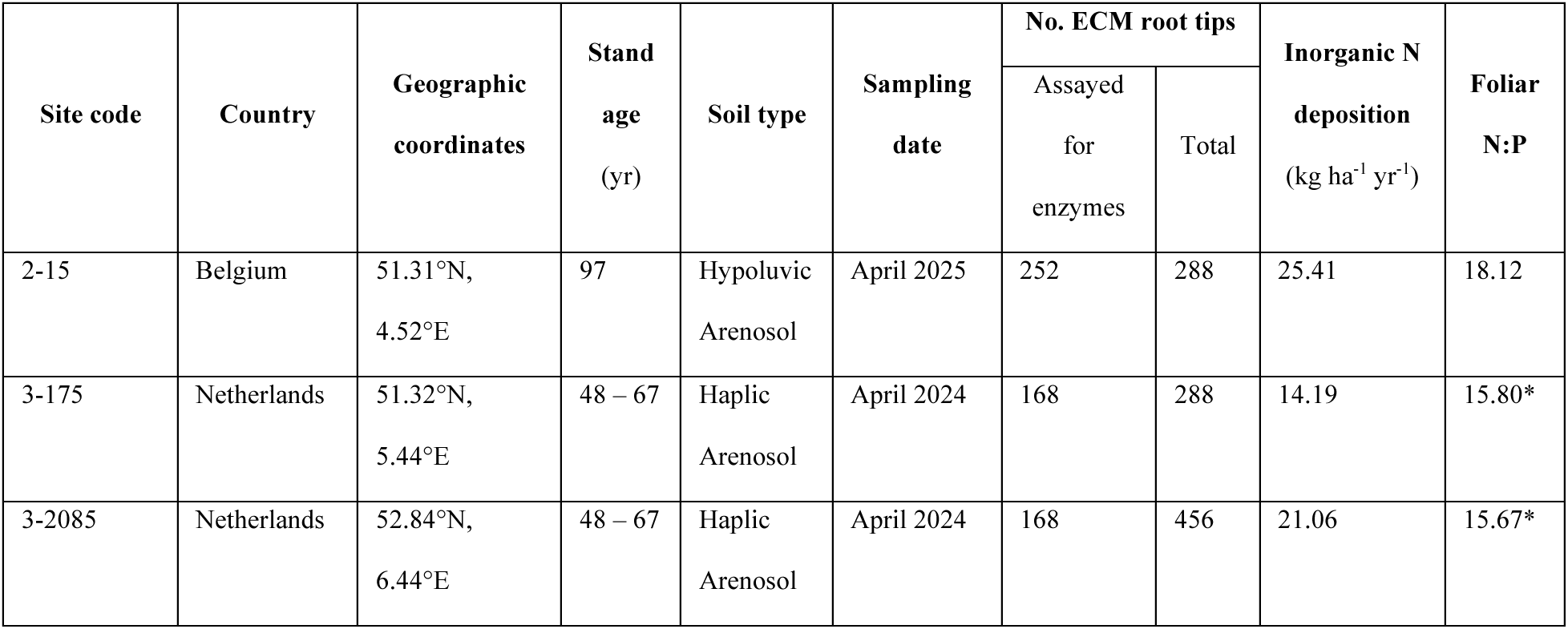
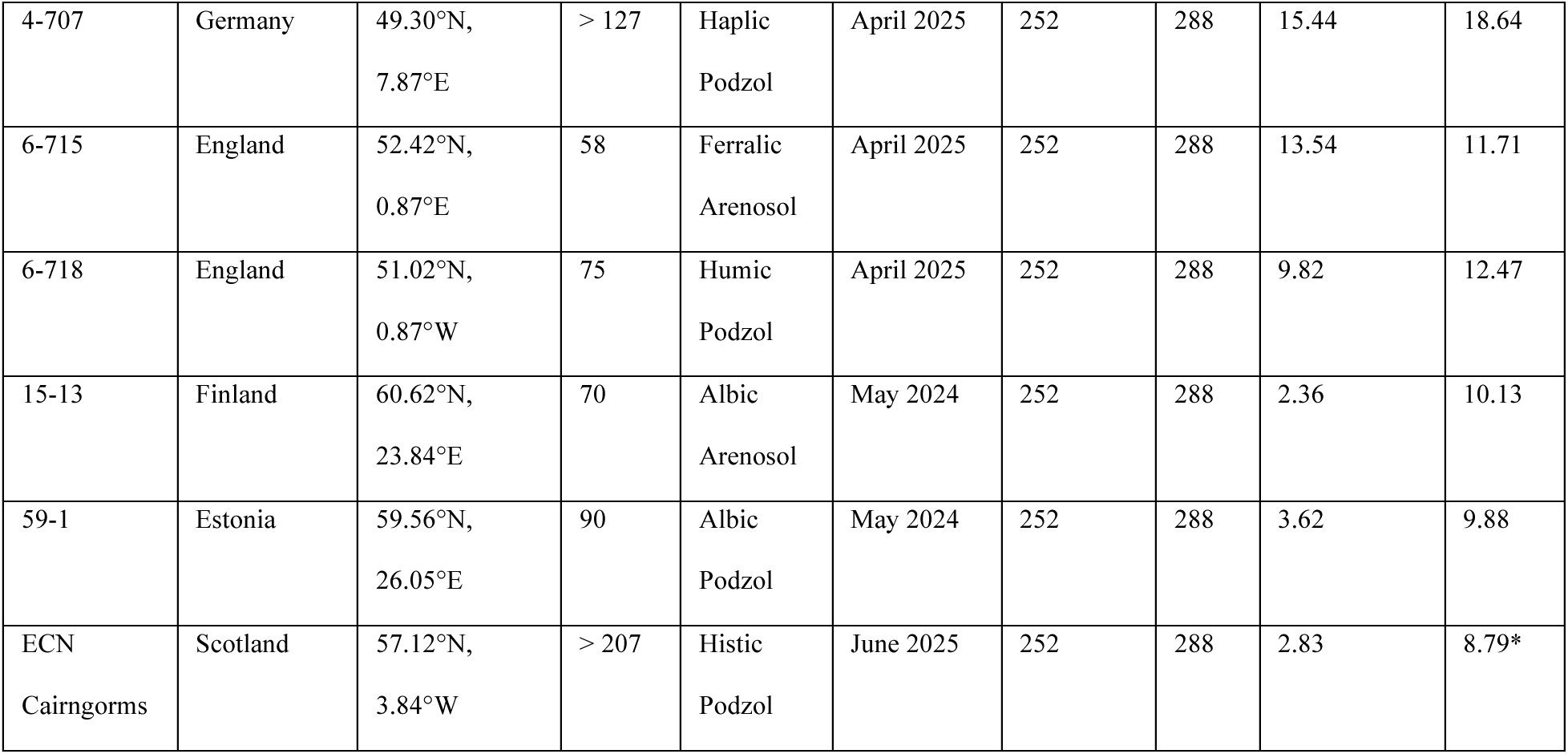
Characteristics of the *Pinus sylvestris* study plots, including N covariates and number of ECM roots collected *versus* assayed for exoenzyme activities. Inorganic N deposition values correspond to total (sum of NH4^+^- and NO3^−^-N) site means averaged over the period 2016–2024 for the ICP Forests plots and 2016-2022 for the SSSI Scottish plot. Foliar N:P ratios marked with an asterisk (*) were unavailable from existing datasets and were therefore measured on leaves collected during sampling.

### Sampling of mycorrhizas

Scots pine roots were collected once per site, from fresh soil samples in the spring of two consecutive years (April-June 2024 and 2025) to avoid potential seasonal bias in ECM exoenzyme activities [50]. At each site, the local ECM community was characterized selecting 288 ectomycorrhizas collected following a standardized protocol [12,13]. Briefly, four evenly spaced soil cores (25 cm deep, 2 cm in diameter) were collected along 24 transects spanning the entire plot, with each transect drawn between a randomly selected tree and its nearest neighbour, while avoiding areas with other ECM host species.

Soil cores were kept sealed in plastic bags and refrigerated at 4°C until processing, which occurred within 10 days. Each soil sample was gently washed through a sieve (0.5 mm mesh size), and three randomly selected live ectomycorrhizas were excised under a dissecting microscope, cleaned, and stored for no longer than 24 h in tap water prior to enzymatic profiling, following [51]. Special care was taken during root selection to exclude fine roots not belonging to the target host tree species. The same set of roots was used for both community taxonomic characterization and enzymatic profiling, except for two sites, where enzymatic assays were performed on additional (site 3-2085) or partial (site 3-175) subsets of roots (**Table 1**).

### Trait measurements

For each ectomycorrhiza, presence-absence of hyphae and rhizomorphs was recorded, and if present were removed prior to the enzymatic assays to enable the standardization of final exoenzyme activity estimates by mantle surface area. The activity of five cell wall-bound exoenzymes was assayed at the level of individual ECM root tips through five sequential incubations in specific fluorometric and colorimetric substrate solutions [52]. In each site, 168 to 252 ectomycorrhizas were assayed (**Table 1**), and negative controls and calibration standards were included.

The selected exoenzymes serve as proxies for N and P acquisition and are hereafter divided into two distinct operational groups [27,53]: (1) nutrient acquisition enzymes, which directly catalyse the release of available N (leucine aminopeptidase, LAP) from proteins and P (acid phosphatase or phosphomonoesterase, PHO) from phosphoesters, and (2) lignocellulolytic enzymes (xylosidase, XYL, acting on hemicellulose, β-glucosidase, GLU, acting on cellulose, and laccase, LAC, involved in the oxidation of phenolic compounds), which indirectly contribute to mobilize N and P by degrading plant and fungal cell walls and necromass, thereby releasing N- and P-containing macromolecules that can subsequently be targeted by nutrient acquisition enzymes. The measured activity of each exoenzyme are assumed to reflect *in situ* expression levels [53], since all assayed enzymes are cell-wall-bound and therefore independent of instantaneous host C supply [25,36].

### Fungal molecular identification

Genomic DNA was extracted from each mycorrhiza using the Extract-N Amp kit (Sigma-Aldrich). The full ITS region was amplified using the primers ITS1F [54] and ITS4 [55]. Amplifications consisted of an initial denaturation at 94°C for 3 min, followed by 35 cycles of 94°C for 30 s, 53°C for 35 s and 72°C for 1 min, with each cycle incrementally extended by 5 s, and a final extension at 72°C for 4 min. PCR products were purified using ExoSap-IT (USB), and Sanger sequenced at Azenta (Oxford, UK). The resulting DNA sequences (.ab1) were basecalled using Phred and assembled using Phrap. Sequences were then trimmed to remove low quality bases and only those of more than 150 base pairs were kept using Trimmomatic. A species hypothesis (SH) was assigned to each sequence using *vsearch* against the UNITE reference database (v. 10FU, 04.04.2024; [56]) at a 97% similarity threshold. When sequences did not match any reference sequence at that threshold, we clustered them together into *de novo* OTUs using *vsearch* and attributed a genus based on the closest sequence in UNITE if above 85% similarity. Remaining sequences were discarded. Ectomycorrhizal status was confirmed by comparing the resulting taxonomic attribution with the FunGuild database [57], and non-ECM taxa were removed.

### Statistical analyses

All statistical analyses were conducted in RStudio (version 2025.05.1+513; R Core Team, 2025). Interspecific variation in enzymatic profiles and functional strategies among dominant ECM fungi (OTUs with ≥ 10 occurrences in the dataset) were visualized using a principal component analysis (PCA) based on species (OTU) mean exoenzyme activities. To link physiology to morphology, species were classified into high- and low-biomass morphological categories [58] according to the presence of rhizomorphs: *Rhizo^+^*when ≥ 10% of individual ectomycorrhizas were observed with rhizomorphs (i.e. high-biomass taxa, grouping medium and long distance soil exploration types according to [59] and [60]), or *Rhizo^−^* (i.e. low-biomass taxa, grouping contact and short distance soil exploration types).

To estimate the dominant strategy in each community, we calculated the community-weighted means (CWMs) enzymatic activity in each plot as:

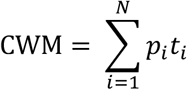

where *p_i_* is the relative abundance of species *i* and *t_i_* is its median enzyme activity. Medians were used to reduce the influence of outliers while not discarding potentially ecologically relevant values. Data distribution was examined for each exoenzyme and normalized using a square-root transformation prior to analyses. The effect of N deposition on enzymatic traits was evaluated for each exoenzyme using linear regressions, considering: a) atmospheric N deposition alone, and b) including stand age as a covariate, since older stands may harbour greater proportions of ECM taxa associated with higher exoenzyme activities reflecting increased reliance on organically bound N and P [61]. No supplementary covariates were added to the models due to the restricted number of independent sites. However, other covariates were expected to have no (e.g. climate) or limited (e.g. soil type) direct effect on ECM exoenzyme activities, with for instance only minor variation in soil categories (**Table 1**) compared to other studies (e.g. [46]). The same models were run in the absence of the Scottish site to observe potential changes in the relationships, as bulk N measurement might be an underestimate of actual throughfall deposition.

The contribution of shifts in community composition along the N deposition gradient and species-level effects were investigated for the two nutrient acquisition enzymes (LAP, PHO) and oxidative capacity (LAC). Sites were partitioned into low and high N deposition categories based on the critical inorganic N load threshold of 5.8 kg ha⁻¹ yr⁻¹ for European ECM communities [13], and exoenzyme means of the five most abundant OTUs were compared between site categories and across all sites. Hereafter, species associated with low and high N deposition refer to species occurring exclusively below and above this threshold, respectively. The contribution of intraspecific enzymatic variation to community-level responses to N deposition was assessed using linear regressions for the same three exoenzymes. Square-root transformed exoenzyme activities were averaged per species and site for widely distributed taxa occurring in ≥ 4 sites with ≥ 3 ectomycorrhizas per site.

Finally, we assessed the potentially buffering effect of ECM community PHO activity on tree N:P ratio. The relationship between ECM community-weighted mean PHO activity and tree N:P ratio was investigated with a Spearman correlation test. Then, assuming that both tree N:P ratio and PHO activity are influenced by N deposition (**Fig. 2**), we quantified the change in the estimated effect (slope) of N deposition on foliar N:P between two models including or excluding community-weighted mean PHO activity.

**Fig. 2.**
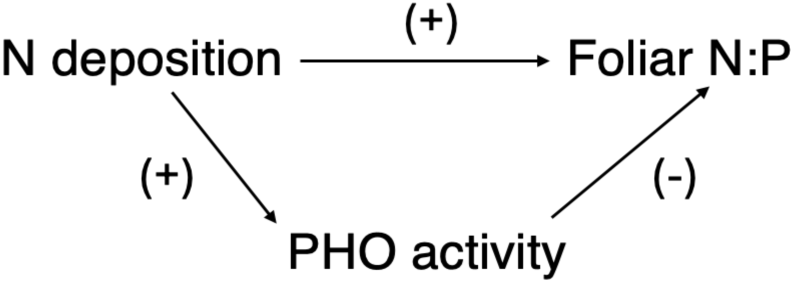
Directed acyclic graph of causal relationships assumed between N deposition, foliar N:P ratio, and ECM community phosphomonoesterase (PHO) activity. N deposition is expected to increase (+) both PHO activity and foliar N:P, while increased PHO activity is expected to decrease (−) foliar N:P ratio.

## Results

### 1. Coordination between morphological and enzymatic traits

Sequences assigned to ECM fungi were obtained for 1 881 roots (155 OTUs) collected across six European countries, among which 1 380 (143 OTUs) had associated enzymatic trait data related to nutrient acquisition (**Supplementary Table 1**).

To summarize the extent of variation in enzymatic traits across species, we performed a principal component analysis for the most abundant species in our dataset (n ≥ 10; **Fig. 3A**). The first two principal components together explain 72.7% of the total variance; PC1 is strongly correlated to hydrolytic enzyme activities, with PHO activity being positively correlated to XYL (*r* = 0.48) and GLU activities (*r* = 0.66), while PC2 contrasts oxidative (LAC) to proteolytic (LAP) activities (*r* = -0.24). Many *Rhizo^+^* species, which include most high-biomass fungi, exhibited high LAP activity (e.g. *Cortinarius aurae*, *Imleria badia*, *Piloderma olivaceum*, *Suillus variegatus*). In contrast, several low-biomass ones, including many species associated with high N deposition (e.g. *Elaphomyces citrinopapillatus, Lactarius subdulcis*) were *Rhizo^−^* and displayed high PHO and lignocellulolytic activities, including oxidative (LAC) activity. On average, selected dominant *Rhizo^+^* taxa exhibited higher LAP activity than dominant *Rhizo^−^*ones, but similar PHO and lower LAC activity (**Supplementary Fig. 1**). Substantial infrageneric variation in enzyme production was detected within most of the six dominant genera in our dataset (**Fig. 3B**), which together account for 71.1% of all mycorrhizas with associated enzymatic trait data. When considering the full observed range, LAP activity varied 91-fold between *Cortinarius pluviorum* and *C. casimiri*, PHO activity varied 13-fold between *Elaphomyces muricatus* and *E. borealis*, and LAC activity varied 27-fold between *Russula ochroleuca* and *R. brunneoviolacea*.

**Fig. 3.**
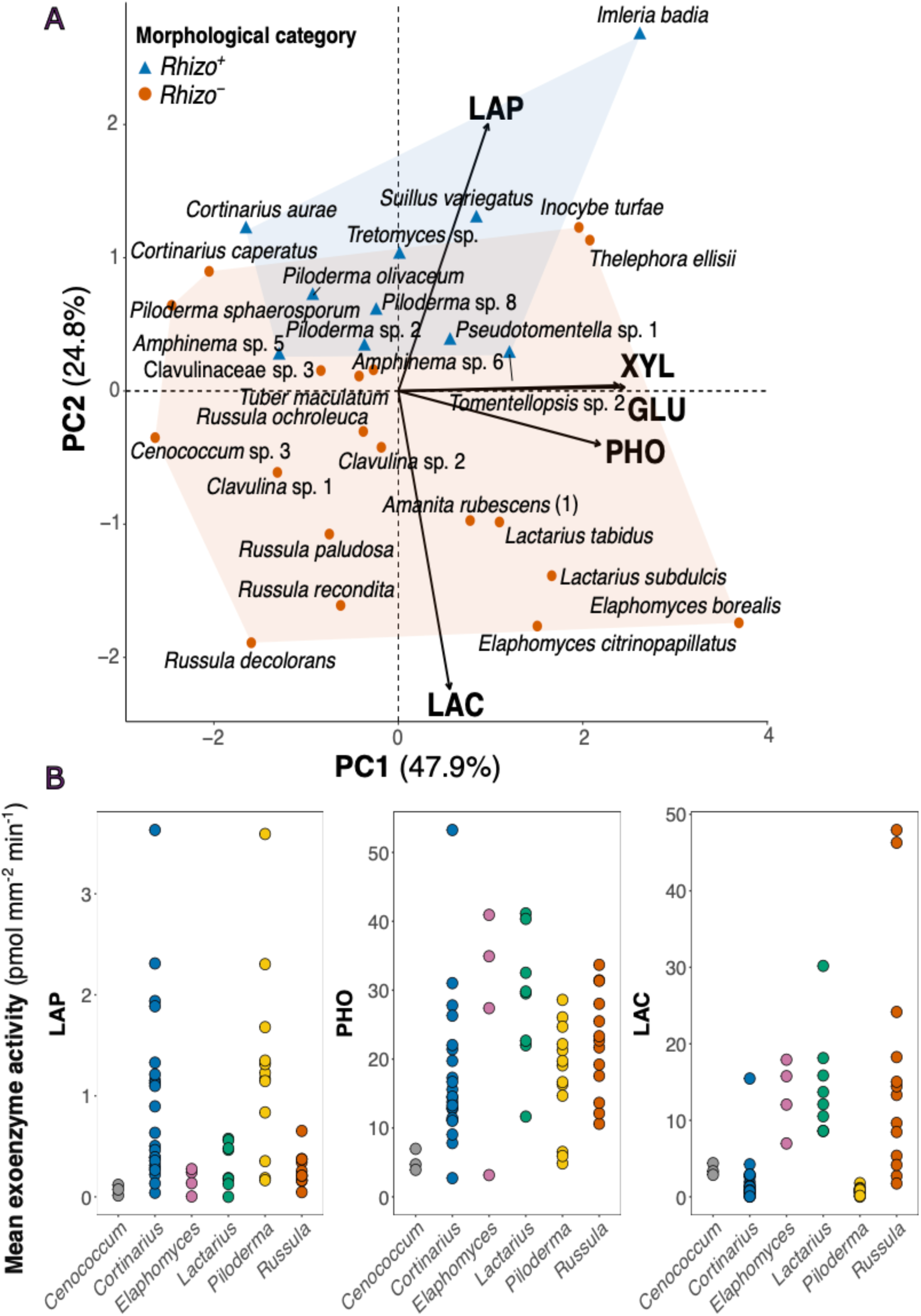
Species-level variation in exoenzyme profiles and morphological traits. **(A)** Principal component analysis (PCA) biplot of five enzymatic traits (square-root transformed), performed on the 29 most abundant species (i.e. OTUs with n ≥ 10). *Rhizo*^+^ (blue triangles and polygon) are the ECM fungal species showing rhizomorphs in at least 10% of ectomycorrhizas, whereas no or less than 10% ectomycorrhizas displayed rhizomorphs in *Rhizo*^−^ taxa (orange dots and polygon). SH numbers have been removed for readability and can be found in **Supplementary Table 1**. **(B)** Variation in exoenzyme production among species within the most dominant genera (with n ≥ 50; i.e. 71.1% for which both OTU assignment and enzymatic trait data were available) for nutrient acquisition hydrolases (LAP, PHO) and one lignocellulolytic oxidase (LAC). Each dot represents one species mean calculated from raw values. Abbreviations: leucine aminopeptidase (LAP), xylosidase (XYL), β-glucosidase (GLU), phosphomonoesterase (PHO), laccase (LAC).

### 2. Atmospheric N deposition correlates with increases in the activity of P-foraging and oxidative enzymes

At the community level, we found decreasing protein N foraging capacity (LAP) with increasing atmospheric N deposition, while all other exoenzymes showed positive increasing trends in relation to N. Significant relationships were detected for PHO (*p* = 0.023) and LAC (*p* = 0.0040) activities (**Fig. 4**). Controlling for stand age and/or excluding the Scottish site strengthened the negative relationship between LAP activity and N deposition, rendering it significant upon exclusion of the Scottish site without stand age adjustment (*p* = 0.045). Conversely, the positive relationship between PHO activity and N deposition appeared more robust to stand age adjustment but only approached significance (*p* < 0.1) in the absence of the Scottish site (**Supplementary Table 2**).

**Fig. 4.**
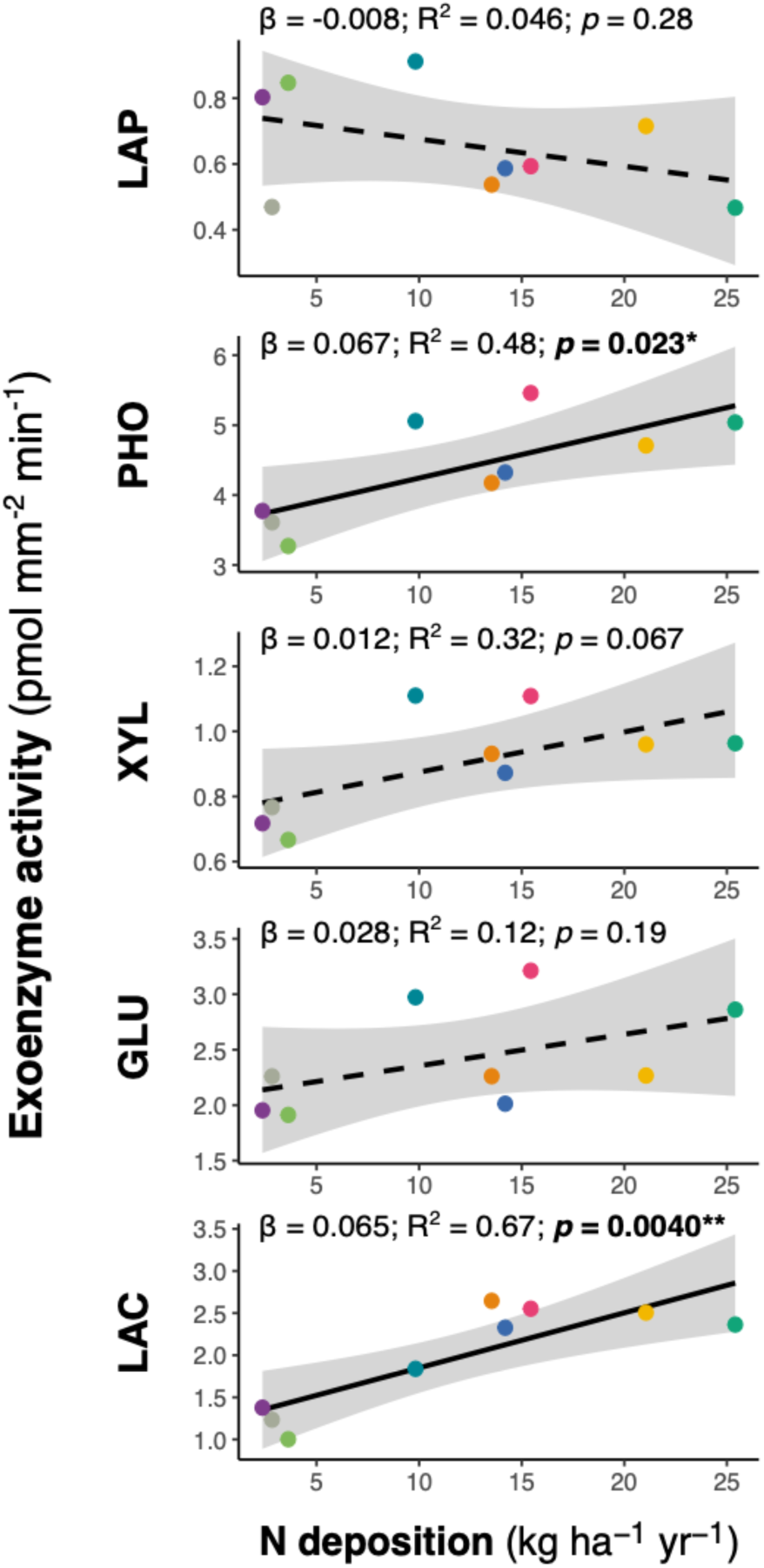
Effect of N deposition on community-weighted mean (CWM) enzymatic traits (square-root transformed). Solid lines indicate significant relationships (*p* < 0.05, linear regressions). The shaded area represents the 95% confidence interval on the mean estimate. Abbreviations: leucine aminopeptidase (LAP), phosphomonoesterase (PHO), xylosidase (XYL), β-glucosidase (GLU), laccase (LAC).

At the species level, some dominant species in low N deposition sites (*Cortinarius caperatus, Piloderma olivaceum*) exhibited higher LAP activity than the overall mean across all species and sites, whereas no dominant species in high N deposition sites did (**Fig. 5**). Conversely, only species from high N deposition sites (*Elaphomyces citrinopapillatus, Lactarius subdulcis, Russula ochroleuca, R. recondita*, together representing 52.1% of all roots across sites receiving > 5.8 kg N ha⁻¹ yr⁻¹) displayed higher PHO activities relative to the mean across all species and sites. The LAC activity patterns were less clearly divided, with dominant species from both high and low N sites showing elevated values and the highest value observed in a low N species (*Russula decolorans*). However, strongly oxidative taxa in high N sites represented a greater proportion (35.4%) of the total communities associated with high N deposition, while *Russula decolorans* accounted for only 4.8% of roots across low N sites.

**Fig. 5.**
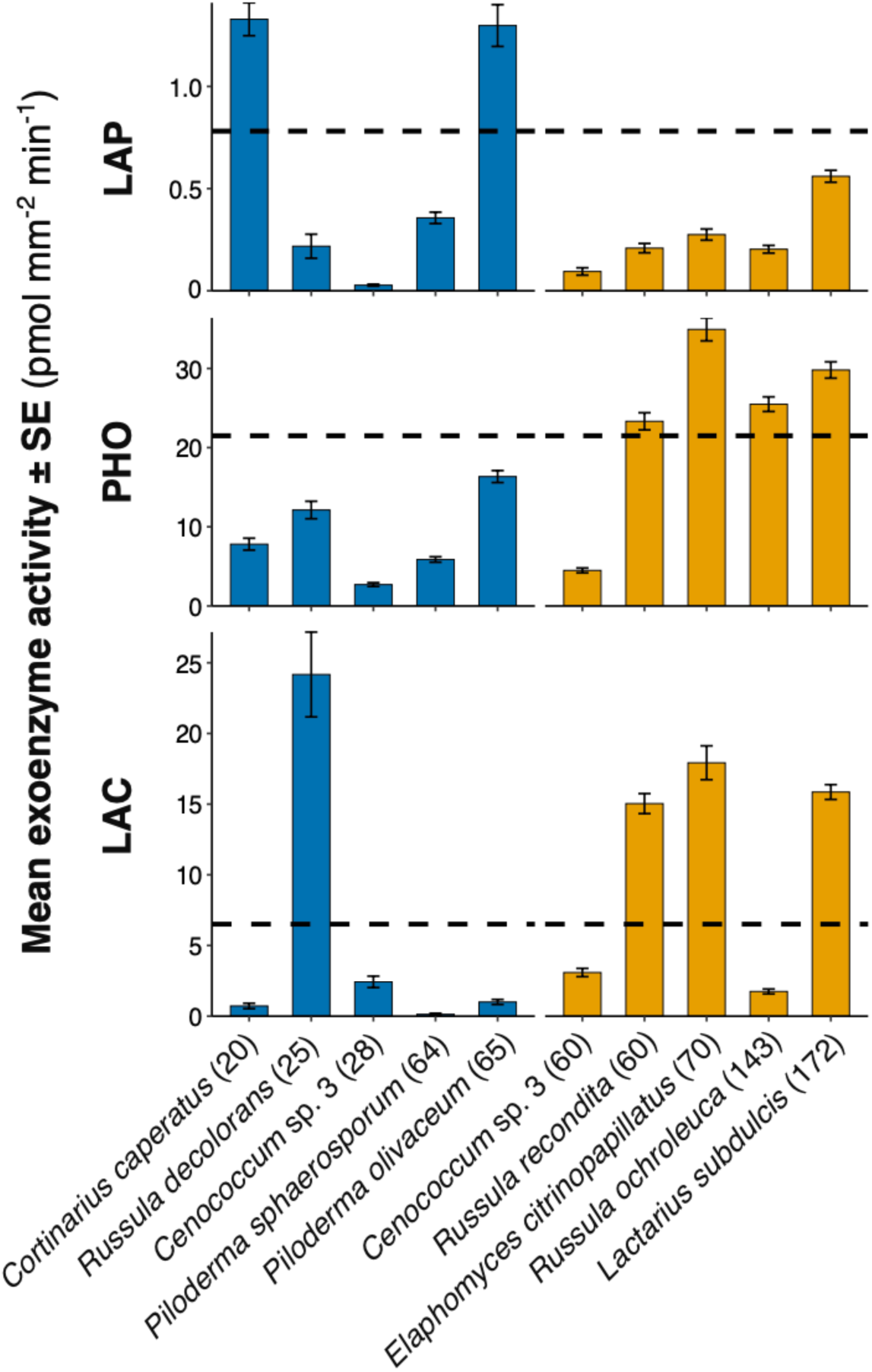
Leucine aminopeptidase (LAP), phosphomonoesterase (PHO) and laccase (LAC) mean activity ± standard error (SE) for the five most abundant ECM species (OTUs) in low (blue bars; left group) and high N sites (orange bars; right group). The selected species together represent 38.4% of the community in low N sites and 59.1% in high N sites. Dashed lines represent the mean for each enzyme across the whole data (all sites combined). Numbers in brackets indicate the number of occurrences of each species in the corresponding site category. SH numbers have been removed for readability and can be found in **Supplementary Table 1**.

No substantial intraspecific enzymatic changes in relation to N deposition were detected in the six selected widespread fungi (**Fig. 6**). Most species showed trends consistent with the community-level changes but none of them were statistically significant, except LAC activity in *Russula paludosa* (*p* = 0.043).

**Fig. 6.**
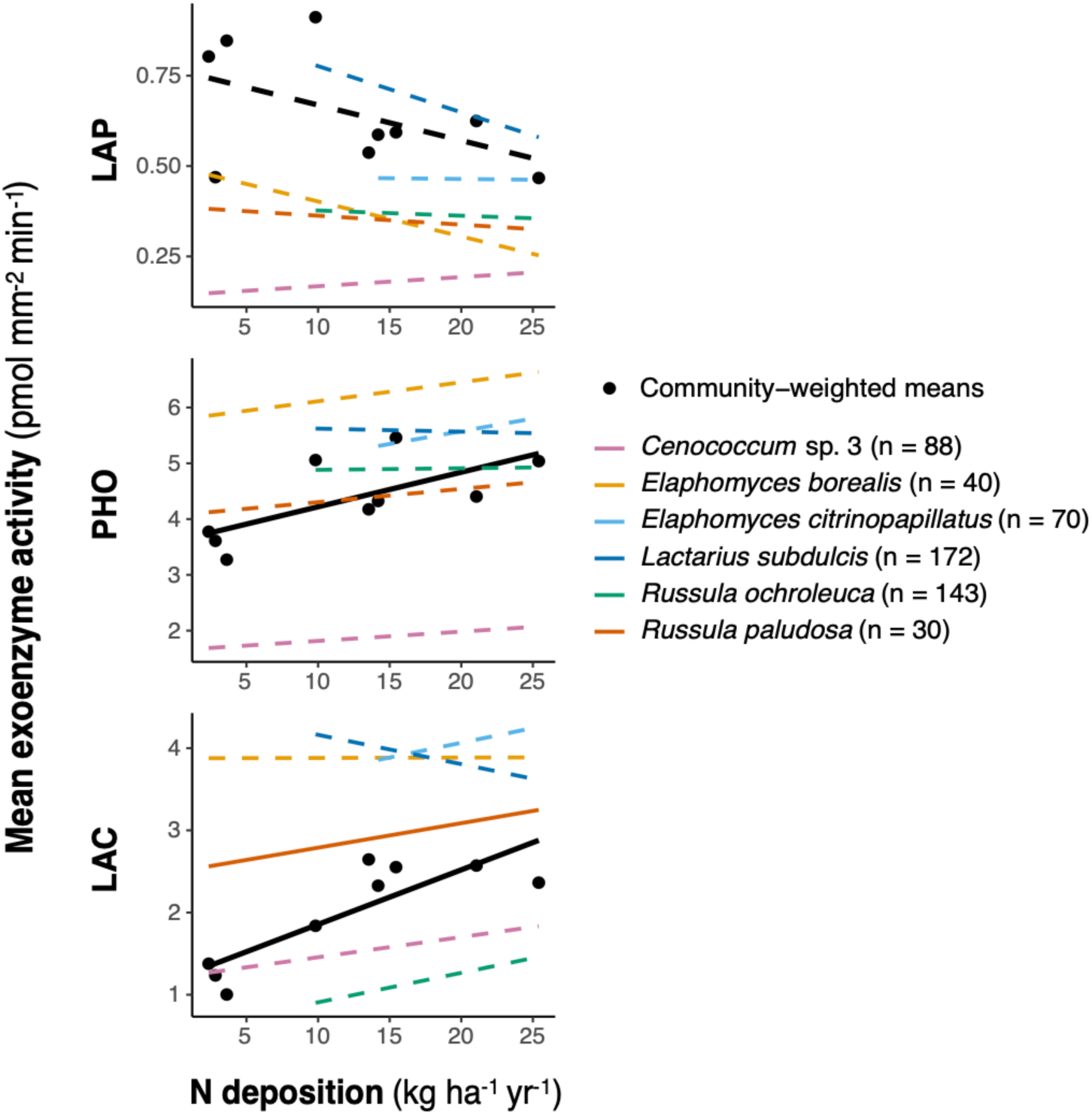
Species trends in leucine aminopeptidase (LAP), phosphomonoesterase (PHO) and laccase (LAC) activities (square-root transformed) for six broadly distributed species along the N deposition gradient. Black dots and lines represent community trends (CWMs). Solid lines indicate significant relationships (*p* < 0.05, linear regressions), while dotted lines indicate non-significant trends. Numbers in brackets indicate the number of occurrences of each species across the whole data. SH numbers have been removed for readability and can be found in **Supplementary Table 1**.

### 3. P-foraging capacity influences tree foliar N:P ratio

A significant positive correlation was found between tree foliar N:P ratio and PHO activity (ρ = 0.87, *p* = 0.0045). Controlling for ECM community PHO activity induced a 37.8% decrease in the estimated effect of N deposition on foliar N:P, pushing the relationship just above the significance threshold (*p* = 0.060; **Fig. 7**).

**Fig. 7.**
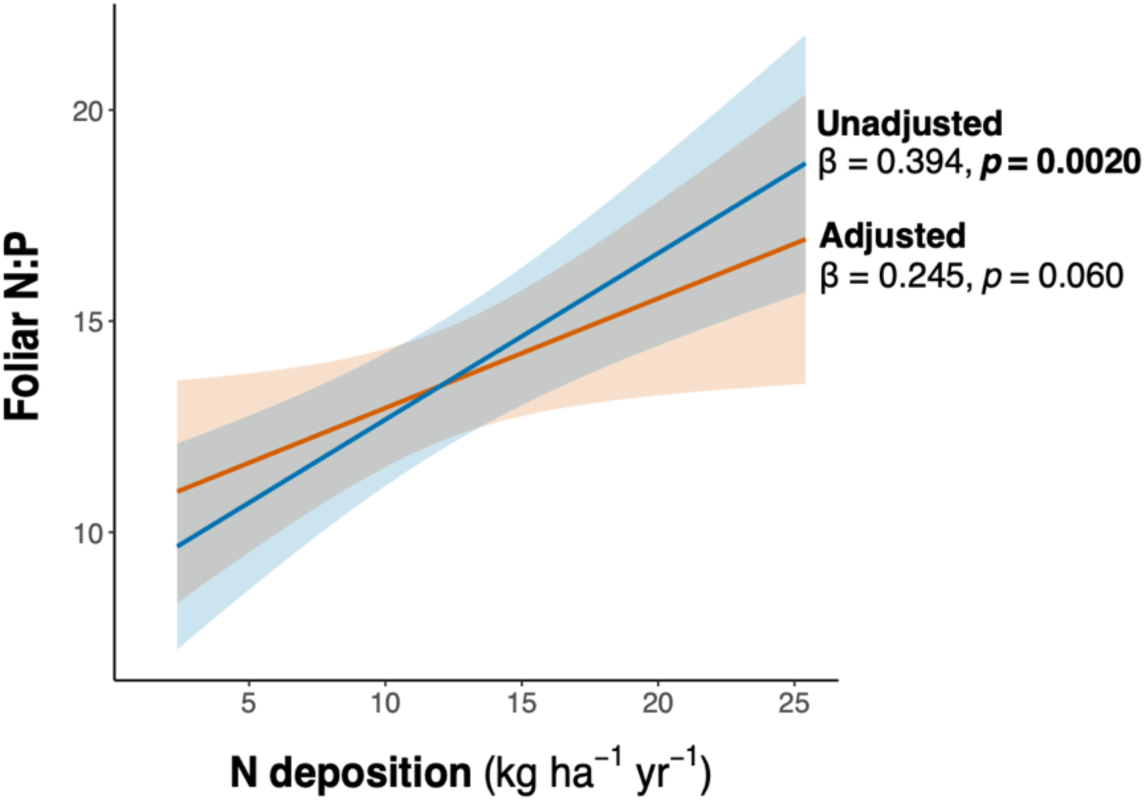
Attenuation of the effect of N deposition on host foliar N:P when controlling for ECM community phosphomonoesterase (PHO) activity. The adjusted model includes community-weighted mean PHO activity as a covariate, which was fixed to its mean value across sites. β and *p*-values refer to the N deposition term in each model.

## Discussion

This study is a rare investigation of the causal mechanisms underlying shifts in ECM community composition along a large-scale atmospheric inorganic N deposition gradient, their links to host tree foliar N:P stoichiometry, and the contribution of intraspecific variation in widespread ECM species to community-level functional shifts. Our results reveal high functional diversity across European ECM assemblages and N-induced changes in community-level enzymatic profiles indicating emerging P limitation (H1). These community-level patterns primarily emerge from pronounced interspecific differences in enzymatic profiles, rather than intraspecific variation (H2), with potential adaptive consequences for host tree performance under anthropogenic nutrient imbalances (H3).

### Community exoenzyme activities indicate N-driven shift towards P limitation

The increase in all community-level exoenzyme activities, except LAP, with increasing N deposition (**Fig. 4**) strongly suggests the emergence of anthropogenic, N-induced P limitation. This is supported by a recent study reporting similar patterns for P-acquiring exoenzymes in Douglas-fir-associated ectomycorrhizas along a regional natural gradient of measured P availability [53], as well as by a documented shift from organic N to P acquisition in ECM communities where N limitation is naturally alleviated, due to having N-fixing hosts [37]. This interpretation is particularly important given that soil P content was not directly measured in the present study, but P limitation was instead inferred from anthropogenic N loads, whose increase does not unconditionally translate into P limitation [20]. For instance, N enrichment has been found to alter tropical ECM communities while suppressing soil phosphatase activity [62]. However, because microbial exoenzyme production was shown to follow ‘economic rules’, with phosphatase activity being enhanced under reduced supply of available P [63], such contradictory results might instead reflect N-induced decline in ECM biomass rather than the emergence of true N-induced P limitation [20].

### Community enzymatic variation primarily reflects species turnover along the gradient

Community-level enzymatic trends were driven mainly by differences in enzymatic profiles among dominant ECM species (**Fig. 3A**), with evidence that these species-specific profiles are locally adapted to the prevailing nutrient limitation (**Fig. 5**), thereby underpinning species distributions along the N-induced P limitation gradient. As previously reported [64], dominant species in communities exposed to chronic N deposition (e.g. *Elaphomyces citrinopapillatus*, *Lactarius subdulcis*, *Russula ochroleuca* and *R. recondita*) exhibited greater potential ability to acquire P from SOM, whereas the opposite was observed across low N sites with some dominant fungi investing more strongly in protein N acquisition (e.g. *Cortinarius caperatus*, *Piloderma olivaceum*). Moreover, the strong positive correlations observed between PHO and lignocellulolytic hydrolases (XYL, GLU), together with the significant increase in LAC activity with N deposition (**Fig. 4**), strengthens the idea that enhanced ECM lignocellulolytic activity may play an indirect but crucial role in alleviating P deficiency [53]. These findings confirm our hypotheses that P limitation is a strong determinant of ECM community structure [65,66], and that N-induced changes in ECM community composition across Europe appear to follow an optimal allocation strategy [47].

While intraspecific response to N deposition generally followed the direction of the community trend, the responses were weak and mostly non-significant (**Fig. 6**). This suggests that intraspecific variation is likely negligible compared to differences between species. Furthermore, only three species (*Cenococcum* sp.*, Elaphomyces borealis, Russula paludosa*) were found in both low and high N sites, suggesting that observed community-level changes remain mostly driven by species turnover. However, the restricted number of sites where these species were present might hamper the generalisation of these results. On the other hand, substantial variation in exoenzyme activity was detected among congeneric species within the most dominant genera (**Fig. 3B**). This could shed light on recent findings documenting contrasting, sometimes opposite, responses of congeneric ECM species to N deposition [13]. This also echoes previous reports of low phylogenetic conservatism in oxidative enzyme production within some ECM genera [67,68], thus indicating that assuming genus-level trait conservatism may be misleading, particularly when enzymatic traits are inferred from taxonomy given the current scarcity and incompleteness of species-level trait data. In addition, the elevated interspecific variation in physiological traits documented here highlights a major limitation in inferring species’ effective function from morphology alone as substantial enzymatic variation was observed within each morphological category, particularly for PHO activity (**Fig. 3A**; **Supplementary Fig. 1**). Despite recent evidence revealing phylogeny-independent genomic signatures of the degradative potential of ECM exploration types [69], our results strengthen the idea of a prevailing effect of evolutionary history on physiology [24], and reinforce the notion that gene content and expressed degradative traits may be substantially decoupled. Importantly, we removed the extraradical mycelium prior to enzymatic assays; thus, our results do not account for the spatial distribution of exoenzymes between the mantle and extraradical mycelium, which may influence morphology–physiology relationships. Regardless, our patterns align with those of [53], who attempted to retain the extraradical mycelium intact for enzymatic analyses, supporting the idea that species identity may exert a stronger influence on enzyme production than morphological categories.

### Efficient P-foragers may partially mitigate host stoichiometric imbalances

We show that the increase in efficient P-foraging ECM fungi partially buffers the effects of N deposition on foliar N:P ratio (**Fig. 7**), confirming that traits of dominant ECM species might regulate ecosystem-level processes [14]. For instance, species such as *Russula ochroleuca* and *Piloderma olivaceum*, which have been previously associated with enhanced plant growth [14,70], are likely promoting tree productivity through distinct nutrient acquisition strategies adapted to the main nutrient limitation (**Fig. 5**). For P, this suggests that trees may continue investing in mycorrhizal-derived P from organic sources as a cost-effective P acquisition strategy under severe N-induced P limitation, contrary to recent findings reporting a shift towards more economical non-mycorrhizal strategies, such as enhanced root release of carboxylates [71,72]. The positive correlation found between foliar N:P and PHO activity might reinforce this view, yet it is most likely explained by the positive effect of N deposition on both foliar N:P and PHO activity. Supporting this hypothesis, it has recently been proposed that plant-derived C allocation to symbiosis is regulated by, and decreases with, the C:N ratio of the ectomycorrhizas themselves [73]. One could expect chronic N deposition to have a decreasing effect on soil and ECM C:N ratios, thereby leading to increased C exudates allocated to efficient microbial P-foragers [74]. This contrasts with previous studies interpreting the success of nitrophilic taxa as the result of a greater tolerance of reduced host C supply under elevated N availability, resulting in uneven communities dominated by a few, less C-costly, competitive taxa [12,13]. Our results instead indicate that these *a priori* low-C-cost taxa, particularly *Rhizo^−^* species (e.g. *Elaphomyces citrinopapillatus*, *Lactarius subdulcis*, *Russula ochroleuca*), exhibit large exoenzyme production that likely entail a non-negligible C cost for the host tree. We therefore suggest that host C allocation to roots and ECM symbionts may be enhanced under low P availability after N limitation is alleviated [20,75]. Under this framework, dominant fungi could be primarily selected for their specialization in organic P mobilization, rather than because of an overall reduced host C cost (i.e. through simplified morphology and physiology).

Our interpretation is further supported by high-biomass C-costly fungi occurring exclusively in high N deposition sites, such as *Scleroderma citrinum* and *Imleria badia*, being associated with elevated PHO activity (**Fig. 3A**; **Supplementary Table 1**). *Imleria badia*, while also exhibiting high proteolytic capacity, has previously been reported to be stimulated under elevated N and low P availability [60,76]. Some high-biomass fungi occurring in low N sites, such as *Suillus* spp., showed elevated PHO activity as well (**Supplementary Table 1**). However, this genus is often considered pioneer, primarily associated with seedlings in early successional stages or recent disturbances [77,78]. Furthermore, a recent study on pine seedlings highlighted elevated species-specific variation among ECM fungi in the coupling between plant C allocation and ECM fungal N exchange [79]. No coupling was observed between N provisioning from *Suillus* and host C allocation, compared to other species (e.g. *Thelephora terrestris*). For P, evidence for reciprocal C-P trade between plant and fungi is generally acknowledged in arbuscular mycorrhizas [80–82] (but see [83]), but it has been little explored in ectomycorrhizas, where interspecific variability needs further characterization [84]. It is however likely that C-P couplings in ectomycorrhizas show elevated variability among species as well, yet mechanisms of host selection based on P are poorly understood [41]. In addition, ectomycorrhizal P-foraging capacity under chronic N inputs is likely driven by fungal N:P disbalances as well, particularly given that fungal N:P was shown to closely match tree foliar N:P [85].

### N-induced increase in ECM oxidative activity could reduce soil C storage

Beyond host tree nutritional status, the consequences for C dynamics of ECM functional shifts in response to N-induced P limitation represent a major unresolved topic [20,86,87], with recent synthesis indicating soil C being positively linked to soil P due to coupled cycling mechanisms [88]. Here, we found N deposition enhanced lignocellulolytic activity, including a significant increase in oxidative activity, with many *Rhizo^−^*species occurring in high N sites displaying both elevated PHO and LAC activities. Added to the N-induced decline of *Rhizo^+^* fungi (**Supplementary Fig. 2**), these morpho-physiological shifts towards dominance by oxidative taxa sequestering little C in their extraradical mycelium indicate a potential ECM-mediated decrease in forest soil C storage in plots exposed to chronic N deposition. However, it should be noted that our study focuses on H_2_O_2_-independent oxidases (e.g. laccase) and does not include H_2_O_2_-dependent peroxidases (e.g. lignin peroxidases; [89]), which might play an important role in ECM nutrient acquisition [27], and whose expression can be suppressed by N deposition [90]. Another study found a suppression of ECM and soil oxidative activity by enhanced N availability [91], which could be explained by methodological discrepancies in the types and categories of oxidases measured. Nevertheless, the strong relationship we found between LAC activity and N deposition remains informative. For instance, many Russulaceae express numerous lignin peroxidase and laccase genes [28,92–95]. Consistent with this, we found high LAC activity in several Russulaceae species (e.g. *Russula* and *Lactarius*, **Fig. 3B**; **Supplementary Table 1**), with many species occurring exclusively in high N sites. Moreover, recent findings showed that ECM fungi identified as oxidative decomposers with elevated Mn-peroxidase production occupy less consistent ecological niches than previously assumed [96] (i.e. old N-limited forest stands [67,68]). Hence, our results still provide evidence that some ECM taxa possessing strong oxidative capacity could be selected under N-induced P limitation and negatively affect soil C dynamics. Interestingly, *Russula ochroleuca*, which is the second most abundant species in our data and is associated with fast tree growth [14], displayed a low PHO:LAC activity ratio (**Supplementary Fig. 3**), meaning its ability to mobilize organic P is not coupled to accelerated soil C mineralization, compared to other high N dominants (e.g. *Elaphomyces citrinopapillatus, Lactarius subdulcis*). This could be of interest in future applied forestry research willing to integrate ectomycorrhizas into sustainable management practices (e.g. tree inoculation with enhanced P-foragers of low oxidative cost).

### Conclusions and perspectives

Our study confirms that species-level enzymatic traits provide a promising toolbox to unravel the mechanisms driving changes in ECM community composition along nutrient availability gradients. Species-specific P-foraging capacities explained species distribution in response to N deposition, with potential adaptive value for host P nutrition under shifting limitations. Future experimental studies should confirm these results under controlled conditions to remove the effect of potential confounding factors on ECM community composition (e.g. climate). Such studies would also clarify the causality between host nutrient status and ECM community composition. Finally, our results could inform applied forestry, notably through the selection of tree mycorrhizal inoculation with enhanced P-foragers of low oxidative capacity to maintain tree nutrient balance with limited impact on soil C storage.

## Acknowledgments

PH was supported by grant 40033782 from the Belgian National Fund for Scientific Research (F.R.S.-FNRS). This project was partly funded by a Kew Pilot Study Grant attributed to GD. We are grateful to the Plant-fungal interaction programme at Kew, grant 11415-101 from Hugh E. M. Osmond through Kew Foundation to LMS and MIB, and to two Kew donors, who wish to remain anonymous. We gratefully acknowledge all representatives of the ICP Forests network (https://www.icp-forests.net/) and NatureScot who provided access to sampling sites and most of the environmental data. Long-term Monitoring of the Cairngorm Environmental Change Network site is supported by the Natural Environment Research Council award number NE /Y006208/1 as part of the NC-UK programme delivering National Capability, and by NatureScot. We thank Sietse van der Linde for his help in the field and valuable advice and Tom Prescott for his help with enzyme measurement.

## Author contributions

GD, PH, LMS and MIB conceived the study. PH, GD, EG, CR and MT collected the data. CA, VA, CL, PM, EV, AV and JW provided access to field sites and environmental data. GD and RG performed bioinformatics, and PH and GD performed data analysis. LMS, GD, TD and MIB obtained funding. PH and GD drafted the manuscript with critical contribution of LMS, MIB and TD. All authors contributed to the revision of the manuscript.

## Data availability statement

The fungal community and trait datasets generated in this study and associated code are available from the authors upon request.

