## Supplementary Fig. 1 for "High nitrogen deposition is associated with phosphorus-efficient ectomycorrhizas in Europe’s Scots pine forests"

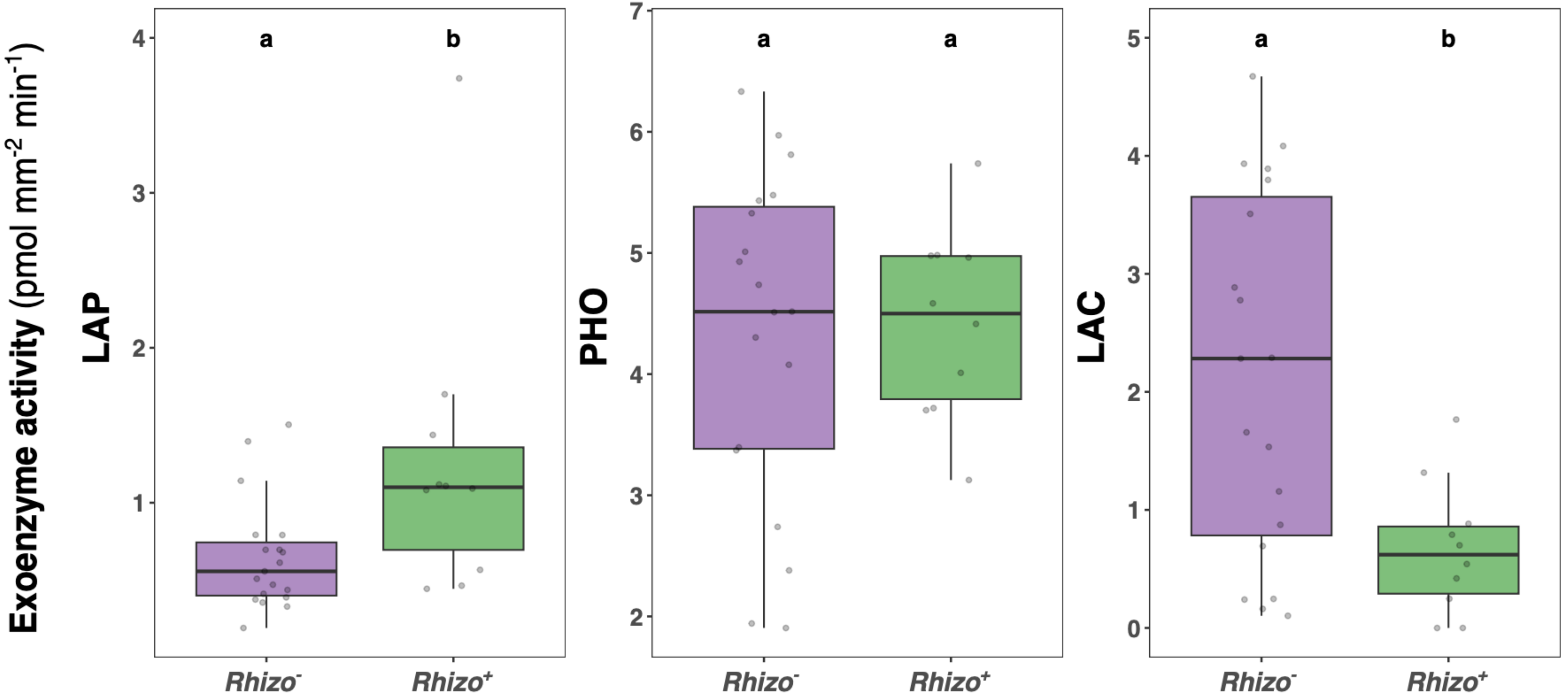


**Supplementary Fig. 1. Leucine aminopeptidase (LAP), phosphomonoesterase (PHO) and laccase (LAC) activities (square-root transformed) in relation to rhizomorph production across the 29 most abundant OTUs ($n \geq 10$).** Different letters indicate a significant difference between morphological groups (Mann-Whitney test, $p < 0.05$).
