## Supplementary Fig. 2 for "High nitrogen deposition is associated with phosphorus-efficient ectomycorrhizas in Europe’s Scots pine forests"

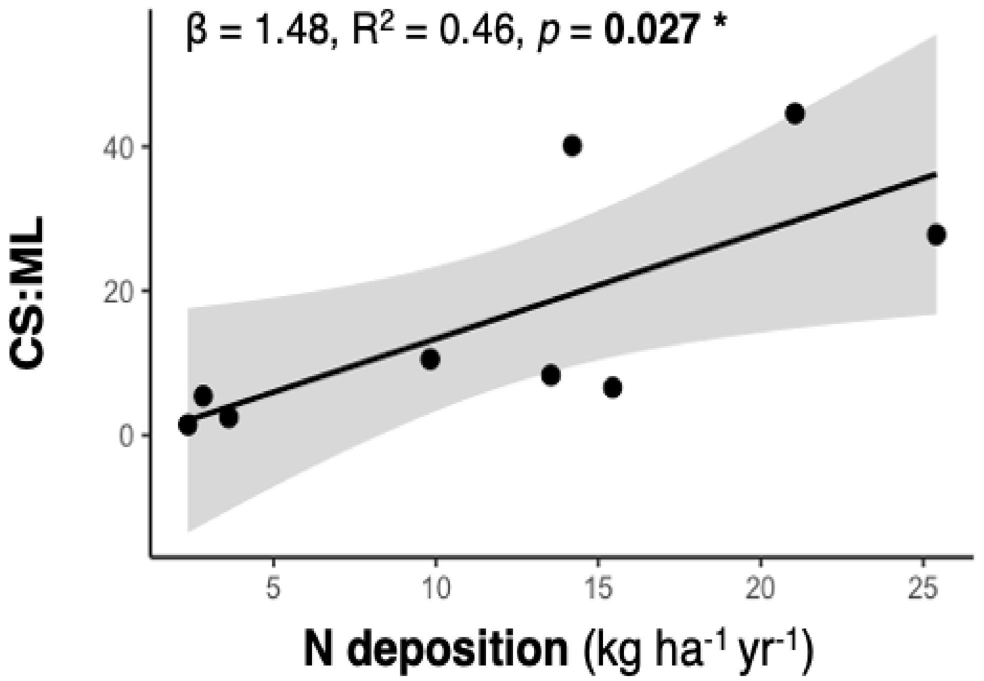


**Supplementary Fig. 2. Effect of N deposition on CS:ML (linear regression).** C, S, M and L respectively correspond to contact, short, medium and long exploration types.
