## Supplementary Fig. 3 for "High nitrogen deposition is associated with phosphorus-efficient ectomycorrhizas in Europe’s Scots pine forests"

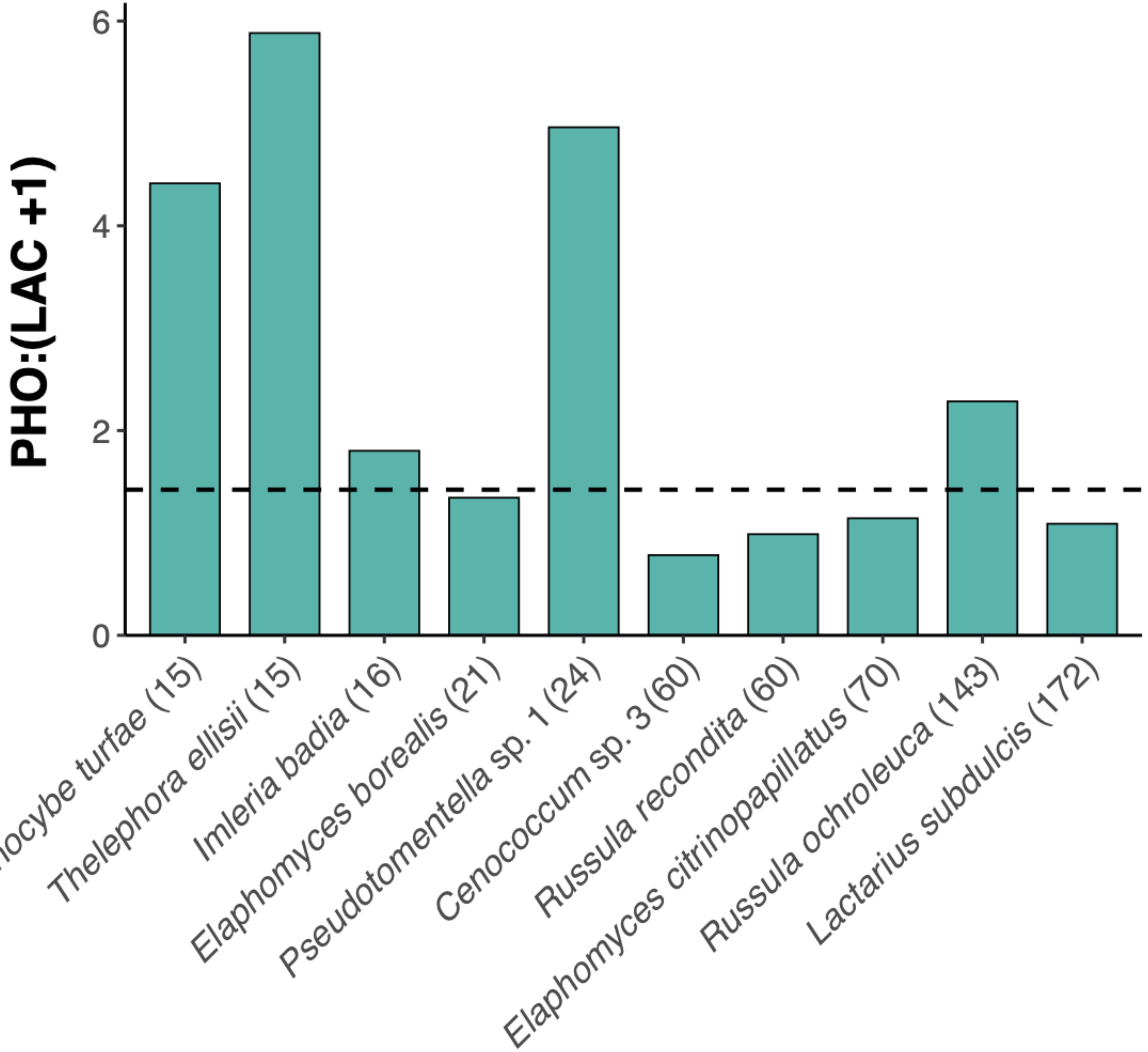


**Supplementary Fig. 3. Phosphomonoesterase (PHO) to laccase (LAC) activity ratio (root-transformed) for the 10 most abundant OTUs in forest sites receiving above ≥ 5.8 kg N ha$^{-1}$ yr$^{-1}$.** Dashed line corresponds to mean ratio value across all these high N sites. Numbers in brackets indicate the number of occurrences of each species across the selected high N sites. SH numbers have been removed for readability and can be found in **Supplementary Table 1**.
